# High quality genome assemblies reveal evolutionary dynamics of repetitive DNA and structural rearrangements in the *Drosophila virilis* sub-group

**DOI:** 10.1101/2023.08.13.553086

**Authors:** Jullien M. Flynn, Yasir H. Ahmed-Braimah, Manyuan Long, Rod A. Wing, Andrew G. Clark

## Abstract

High-quality genome assemblies across a range of non-traditional model organisms can accelerate the discovery of novel aspects of genome evolution. The *Drosophila virilis* group has several attributes that distinguish it from more highly studied species in the *Drosophila* genus, such as an unusual abundance of repetitive elements and extensive karyotype evolution, in addition to being an attractive model for speciation genetics. Here we used long-read sequencing to assemble five genomes of three virilis group species and characterized sequence and structural divergence and repetitive DNA evolution. We find that our contiguous genome assemblies allow characterization of chromosomal arrangements with ease and can facilitate analysis of inversion breakpoints. We also leverage a small panel of resequenced strains to explore the genomic pattern of divergence and polymorphism in this species and show that known demographic histories largely predicts the extent of genome-wide segregating polymorphism. We further find that a neo-X chromosome in *D. americana* displays X-like levels of nucleotide diversity. We also found that unusual repetitive elements were responsible for much of the divergence in genome composition among species. Helitron-derived tandem repeats tripled in abundance on the Y chromosome in *D. americana* compared to *D. novamexicana*, accounting for most of the difference in repeat content between these sister species. Repeats with characteristics of both transposable elements and satellite DNAs expanded by three-fold, mostly in euchromatin, in both *D. americana* and *D. novamexicana* compared to *D. virilis*. Our results represent a major advance in our understanding of genome biology in this emerging model clade.

**Significance statement:** The *Drosophila virilis* sub-group is an emerging model with an enticing combination of attributes, including abundant and diverse repetitive DNA content, structural rearrangements, and hybridization capability. The lack of high-quality genome assemblies for this group have prevented detailed understanding of genome evolution. Here, we present five new long-read genome assemblies of three virilis sub-group species along with analyses of structural variants, polymorphisms, repetitive DNAs, and Y chromosome genes and repeats. We find that the expansion and mobilization of non-canonical repetitive elements accounts for most of the divergence in assembled genome sequence between these species, with an especially striking takeover of the Y chromosome by a single type of element in one of the three species. Overall, our study positions the virilis sub-group as a model for a variety of future studies.

---

Recent advances in genome sequencing technologies have made it possible to generate high quality assemblies of non-model organisms with relative ease (Ellegren 2014; Kim *et al*. 2021). Coupled with population resequencing, these advances can now accelerate discoveries in comparative genomics research and reveal previously recalcitrant attributes of genome evolution, such as repetitive DNA and heterochromatin. This promise is particularly prevalent among insect species that represent “neomodel” organisms, particularly Dipterans that have relatively small genome sizes and can serve as laboratory model systems. In addition, the progress in genome editing technologies necessitates the availability of high quality annotated genomes to maximize the utilization of these neo-model organisms for a variety of evolutionary and molecular genetic studies (Dickinson *et al*. 2020).

The *Drosophila virilis* species group is an emerging model for comparative genomics and genome evolution. The virilis sub-group (*D. americana, D. novamexicana, D. lummei*, and *D. virilis*) originated in temperate forests in Eurasia ∼8.5 million years ago (mya) after splitting from the *D. littoralis* sub-group and subsequently colonized other temperate regions in northern Eurasia and North America in concurrent speciation events that took place ∼5 mya (*D. virilis* split), ∼3 mya (*D. lummei* split), and ∼1 mya (*D. americana/D. novamexicana* split; Figure 1) (Throckmorton 1982; Yusuf *et al*. 2022). They are thought to inhabit slime flux in riparian habitats and can be found in a variety of warm/humid habitats in marshy regions, particularly in North America (Blight and Romano 1953). Recent work has highlighted this species group’s utility in a variety of research programs, including neurobiology and behavior (LaRue *et al*. 2015), genome evolution (Flynn *et al*. 2020b, 2023a; Fonseca *et al*. 2013), pigmentation evolution (Wittkopp *et al*. 2009; Sramkoski *et al*. 2020; Ahmed-Braimah and Sweigart 2015), and speciation (Sweigart 2010a,b; Ahmed-Braimah 2016). We specifically emphasize its utility in comparative genomics and evolutionary genetics research programs because hybrids between all species in this group can be generated easily in the laboratory, which makes many experimental approaches tractable. For example, this species group is particularly suited for research on the genetic basis of reproductive isolation because most hybrid crosses produce a small percentage of fully viable and fertile progeny that can be used for genetic mapping or genomic introgression analyses (Sweigart 2010a,b; Ahmed-Braimah and Sweigart 2015; Ahmed-Braimah 2016).

**Figure 1.**
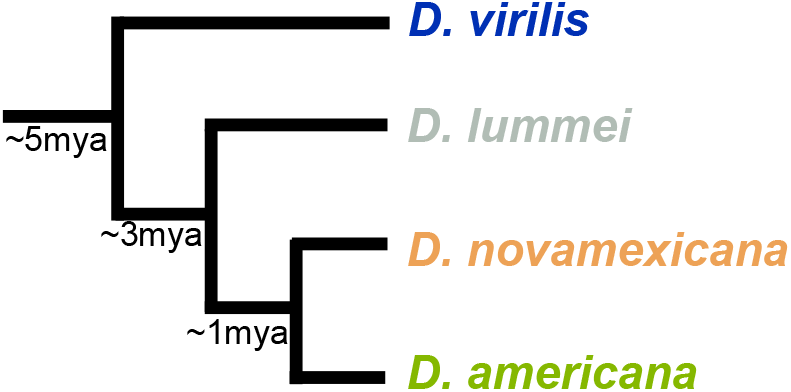
Phylogenetic relationships of virilis sub-group members and divergence times.

**Figure 2.**
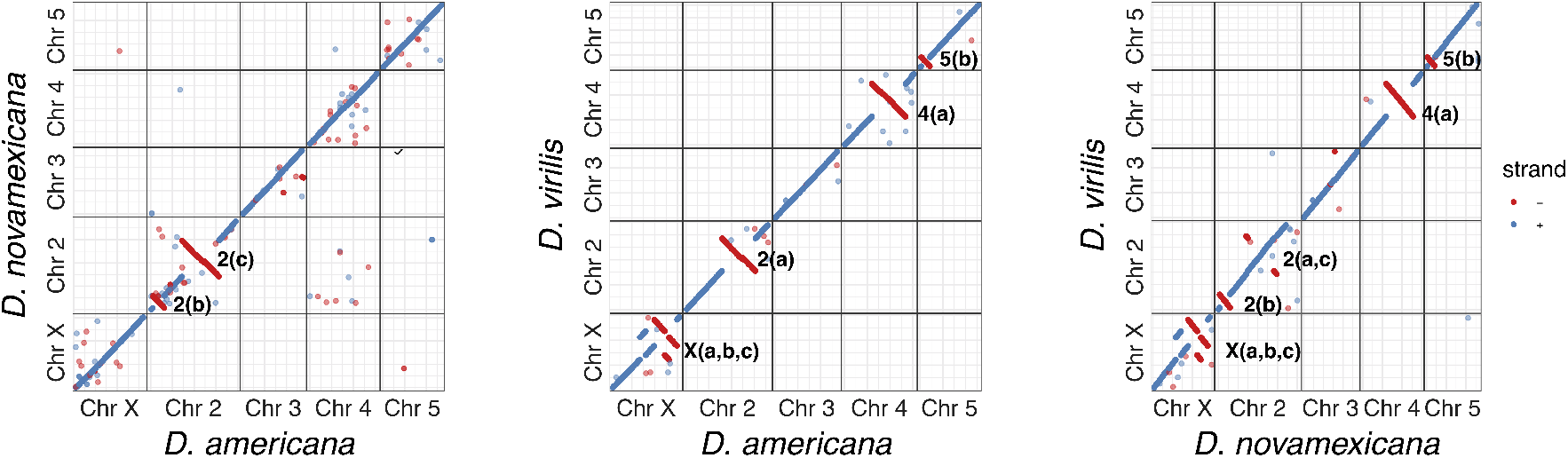
Pair-wise whole chromosome alignment dotplots between the scaffolded PacBio genomes of the three species. The identified inversions are indicated on the right of the negative strand notation of each inversion.

The virilis sub-group has also served as a model for karyotype evolution and chromosomal rearrangements (Reis *et al*. 2018). This species group maintains the ancestral *Drosophila* genus configuration of Muller element karyotypes, but has undergone several chromosomal fusions and structural rearrangements between members (McAllister 2003; Caletka and McAllister 2004). Early work has identified a number of species-specific and/or shared chromosomal inversions, thus providing a template to investigate the causes of chromosomal inversions and their evolutionary consequences (Reis *et al*. 2018). One member of the virilis sub-group, *D. americana*, has evolved two chromosomal fusions—2-3 and X-4—while all remaining members of the virilis group maintain the ancestral karyotype (Hughes 1939). Furthermore, the X-4 fusion in *D. americana* is clinally distributed, suggesting a functional role for the fusion as a mechanism to influence chromosome-wide spatial allele frequencies (McAllister 2002).

The *D. virilis* genome is rich in repeats, most notably 7 bp satellites that compose 40% of the genome, most of which are not included in the genome assemblies produced for this species (Flynn *et al*. 2020b; Clark *et al*. 2007). Transposable elements also occupy a high proportion of the virilis genome, but their genome-wide evolution and turnover among species has not been investigated. Repeats are generally classified as either interspersed (IR) or tandem (TR), having distinct localization patterns and mechanisms of copy number change (Biscotti *et al*. 2015). IRs are typically transposable elements that are interspersed because of their ability to mobilize, and TRs are typically satellite DNAs, which form long continuous arrays that are enriched near centromeres and telomeres. *D. virilis* contains multiple abundant elements that cannot be neatly categorized because they occur in medium length tandem arrays that are interspersed in multiple locations across the genome (Heikkinen *et al*. 1995; Abdurashitov *et al*. 2013; Silva *et al*. 2019; Kuhn *et al*. 2021). How such interspersed-and-tandem repeats—which we will call ITRs—evolve is not known. Studying their abundance and distribution in the genomes of closely related species will reveal whether they mainly evolve by moving to new locations (like IRs) or by altering the length of their arrays (like TRs).

Little is known about the evolutionary dynamics of the Y chromosome among closely related species, owing in part to the difficulty of reliably assembling contiguous pieces of this male-limited, highly repetitive chromosome in most genome assembly efforts. The Y chromosome in Drosophila carries essential male fertility factors, and can thus be a major source of cross-species incompatibilities (Lamnissou *et al*. 1996; Vigneault and Zouros 1986; Heikkinen and Lumme 1998). Drosophila Y chromosomes experience less constraint than the rest of the genome because they are almost completely silenced in a heterochromatic state. Having a lower effective population size than the rest of the genome and lacking recombination provides an opportunity for selfish repetitive elements to proliferate rapidly on the Y chromosome, potentially leading to species-specific Y chromosome repetitive DNA composition, which may contribute to hybrid incompatibilities between species. Indeed, the *D. novamexicana* Y chromosome is necessary and sufficient to cause hybrid male sterility in *D. novamexicana*/*D. virilis* hybrids (Heikkinen and Lumme 1998). Understanding the contributions of the Y chromosome to speciation requires improvements in genome assembly and analytical tools to annotate and analyze its content (Chang *et al*. 2022).

Here we generated high-quality genome assemblies for three strains of *D. americana* and one strain each of *D. novamexicana* and *D. virilis* using Pacific Biosciences or Oxford Nanopore long-read sequencing technologies. For each genome we used high molecular weight DNA from males to specifically assemble Y chromosome contigs. We first show that breakpoints of fixed inversion differences between species and within *D. americana* are easily identified to nearly single base pair resolution upon pair-wise whole chromosome alignment. Furthermore, we are able to confidently identify the highly repetitive Y chromosome-derived contigs for each assembly, for which we found transient protein-coding gene content and repetitive DNA turnover. Finally, we describe pervasive patterns of transposable element and repetitive DNA evolution among this trio of species. Our results represent a major advance in our understanding of genome evolution among this group of species and provide excellent resources for additional comparative genomics research in this emerging model clade.

## Results and Discussion

### Assembly attributes

We generated several assemblies for the three species using at least one of three assembly programs: Canu (Koren *et al*. 2017), Falcon (Chin *et al*. 2016), and MECAT (Xiao *et al*. 2017). The three assemblers achieved varying assembly sizes and contig N50 values (Table 1). Genome size in *D. virilis* has previously been reported to be ∼400 Mb according to flow cytometry estimates (Bosco *et al*. 2007), however sequence assembly sizes often fall short of that. Here we find that the largest assembly among the three species is ∼316 Mb for *D. americana*-G96, with the remaining assemblies being in the range of 189–283 Mb (Table 1). This discrepancy in genome size between flow cytometric measures and sequence assemblies is largely due to the underrepresentation of highly repetitive stretches of 7 bp-unit length satellite DNA that characterizes this species group (Flynn *et al*. 2020a). The most contiguous assemblies were achieved with Canu for the two *D. americana* Nanopore genomes, with N50’s comparable to the average Muller element size, while only the scaffolded PacBio assemblies reached comparable N50 values. Thus, these assemblies represent a notable advance in assembly quality and contiguity for this trio of species.

**Table 1.** Assembly details.

| Species | Seq. tech. | Assembly* | Total length | # of contigs/scaffolds | Mean length | contig/scaffold N50 |
| --- | --- | --- | --- | --- | --- | --- |
| *D. americana* (G96) | PacBio | Canu | 316,316,524 | 530 | 448,889.43 | 1,842,965 |
| *D. americana* (G96) | PacBio | Falcon | 283,249,233 | 631 | 596,823.63 | 2,807,137 |
| *D. americana* (G96) | PacBio | Quickmerge (s) | 214,229,691 | 286 | 749,054.86 | 27,942,935 |
| *D. americana* (ML97.5) | Nanopore | Canu | 236,071,707 | 322 | 733,141.95 | 28,327,896 |
| *D. americana* (SB02.06) | Nanopore | Canu | 240,772,203 | 465 | 517,789.68 | 31,320,009 |
| *D. novamexicana* | PacBio | Mecat | 182,394,971 | 292 | 921,186.72 | 3,158,326 |
| *D. novamexicana* | PacBio | Mecat (s) | 182,394,971 | 198 | 624,640.31 | 30,194,120 |
| *D. virilis* | PacBio | Mecat | 189,443,829 | 222 | 853,350.58 | 8,697,263 |
| *D. virilis* | PacBio | Mecat (s) | 189,443,429 | 185 | 1,024,018.54 | 29,144,718 |

### Structural rearrangements

The virilis sub-group is an excellent model for karyotype evolution because of rapid changes in whole chromosome fusions and chromosome rearrangements (Hughes 1939; Reis *et al*. 2018; Flynn *et al*. 2023b), however the mechanisms that facilitate these mutational events are not well understood and require characterization of sequence elements that facilitate rearrangement events. To examine the dynamics of inversions between the three species genomes we first identified the major inversion differences using whole-genome alignments of the scaffolded PacBio assemblies. We observed several of the known inversion differences between species (Reis *et al*. 2018), and were able to pinpoint inversion breakpoints to ≤1 kb resolution (File S1). *D. americana*-G96 and *D. novamexicana* contain two inversion differences on chromosome 2: *In(2)a* and *In(2)b*, while *D. virilis* contains several fixed inversion differences compared to *D. americana* and *D. novamexicana* in all chromosomes except chromosome 3, which is collinear between the three species. The X chromosome carries the most fixed inversions between *D. virilis* and the other two species, with several nested inversions on the centromere proximal region of the chromosome (Figure 3). Among the three *D. americana* strains we find that the second chromosome inversions are fixed, but several inversions on the other chromosomes are polymorphic, including *In(4)a, In(5)a, In(5)b, In(X)b and In(X)c* (Figure S1).

**Figure 3.**
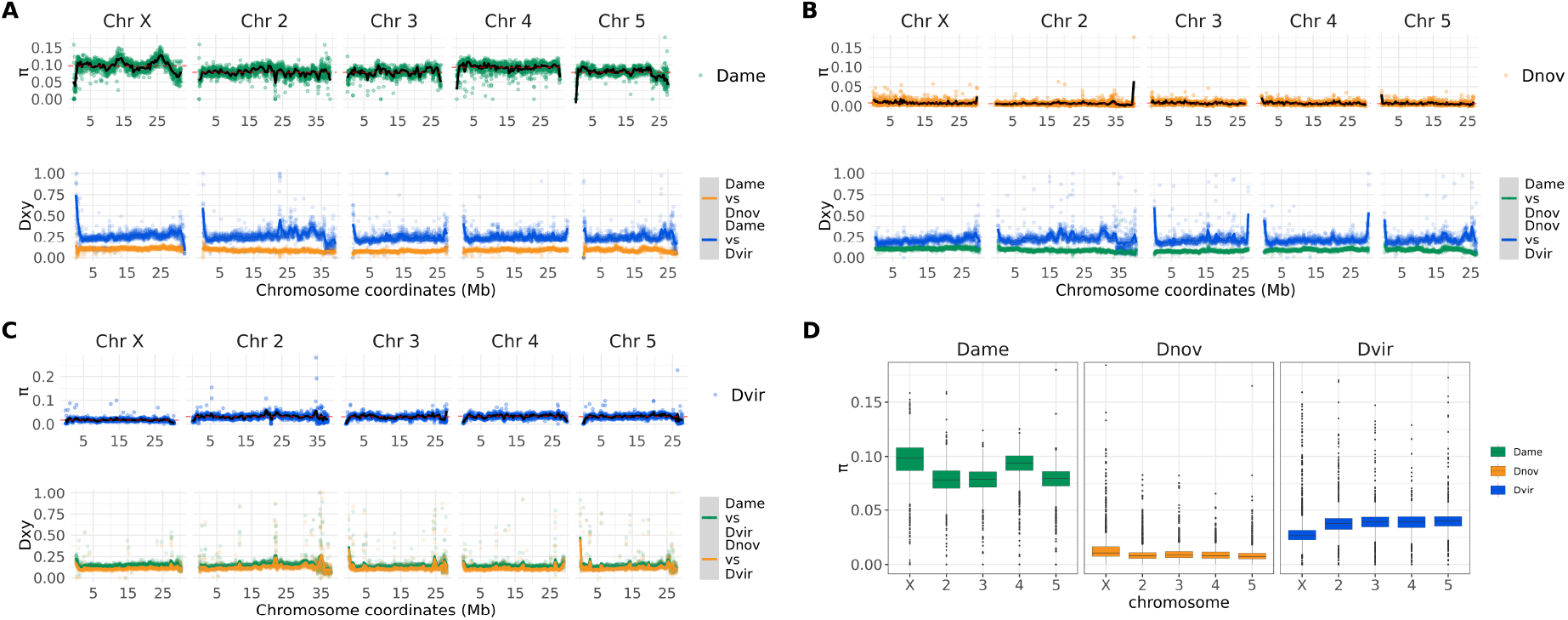
Patterns of nucleotide diversity and divergence in the virilis group. Intraspecific nucleotide diversity (*p*) is plotted along the five major chromosome arms for *D. americana* (A; top), *D. novamexicana* (B; top), and *D. virilis* (C; top), while pairwise divergence (*D*_*xy*_) is plotted at the bottom of each panel figure of the focal species. (D) Box plots of nucleotide diversity for each major chromosome.

Previous work has found that DAIBAM MITE (miniature inverted-repeat TEs) are enriched and potentially causal near the inversion breakpoints (Evans *et al*. 2007; Reis *et al*. 2018). We found that these DAIBAM elements were indeed enriched in breakpoint regions within *D. novamexicana* compared to the rest of the genome with a permutation test (*p* < 0.001). However, these elements were mostly found in *D. novamexicana*, and were never found in the same homologous breakpoint region across all three species, and rarely across only two species (Figure S2). Rapid evolution of repetitive sequences makes it difficult to identify whether DAIBAM accumulation occurred before or after the inversions, and thus it is difficult to assess whether this element is causal.

### Divergence and polymorphism

We examined the pattern of divergence and polymorphism among the three species using shortread sequence data from population samples (Table S1; Ahmed-Braimah *et al*. (2017); Flynn *et al*. (2020a)). We calculated intraspecific nucleotide diversity (*p*) and interspecific pairwise divergence (*D*_*xy*_) in 20 kb non-overlapping windows across the five major chromosomes (Figure 3). Our results show that *D. americana* has the highest average nucleotide diversity (*p*=0.08), while *D. novamexicana* and *D. virilis* have lower average nucleotide diversity (*p*=0.007 and *p*=0.02, respectively). These results are consistent with demographic histories and biogeographic distributions among these species (McAllister 2002; Mirol *et al*. 2008; Fonseca *et al*. 2013). *D. americana* has maintained a large effective population size and occupies a wide habitat, while *D. novamexicana* is restricted to a narrower range west of the Rocky mountains (Caletka and McAllister 2004). The small effective population size of *D. novamexicana* likely contributes to the low level of nucleotide diversity within this species—it is worth noting, however, that strain sampling of these species has historically been poor. *D. virilis*, on the other hand, has recently undergone a range expansion that resulted in the species’ colonization of multiple continents outside of its ancestral range in Eurasia (Mirol *et al*. 2008). Thus, its global populations have experienced multiple founder events that likely contributed to reduced genetic variation.

Nucleotide diversity also differed between the autosomes and sex chromosomes among the three species. Specifically, the X chromosome showed elevated nucleotide diversity in *D. americana* and *D. novamexicana* compared to the autosomes, but reduced nucleotide diversity in *D. virilis* compared to the autosomes (*p* < 0.00001, Wilcoxon Rank Sum Test; Figure 3D). Interestingly, the fourth chromosome of *D. americana*—which is fused to the X in northern populations of *D. americana*—shows elevated nucleotide diversity that is similar to the X chromosome (*p* < 0.00001, Wilcoxon Rank Sum Test; Figure 3D). These results beg the question of how the three species’ evolutionary histories resulted in differing patterns of nucleotide diversity on the autosomes and X chromosome. The X-4 fusion in *D. americana* is clinally distributed, with the fused variant being at high frequency in northern populations and the unfused variant at high frequency in the southern range. Here we use a collection of strains from throughout this range, which likely explains the higher levels of nucleotide diversity between distinct northern and southern haplotypes. Because chromosome 4 is fused to the X chromosome, it effectively behaves as a neo-Y chromosome in southern populations where the fused X-4 genotype is absent. Thus, nucleotide diversity measurements within our diverse samples is elevated for the X and fourth chromosomes relative to the remaining autosomes.

Our analysis of divergence between the three species’ genomes shows that divergence patterns across the genome are largely invariant at this analysis resolution, but a few genomic windows show elevated divergence (Figure 3). The overall divergence pattern reflects species relationships, and the autosomes show a similar pattern of divergence to the X chromosome. Notably, only some of the centromeric/telomeric ends of the chromosomes show elevated divergence, potentially indicating the limit of the assembly into centromeric heterochromatin.

### Identification of Y chromosome contigs and protein-coding genes

Genome assemblies typically lack contig assignment to the Y chromosome, owing to the difficulty in identifying these contigs. Here we sought to assign Y chromosome origin to contigs by using information derived from male and female short-read DNA sequence data. Y-derived contigs are expected to exhibit skewed coverage ratios between males and females. Specifically, the coverage ratio (female/male) for Y contigs should be ∼0, while the coverage ratio for autosomes and the X chromosome should be ∼1 and ∼2, respectively. We calculated the average coverage from mapped male and female reads for each contig and found that, indeed, contig coverage ratios are distributed according to the expectation based on their chromosomal origin (Figure S3A). We also used a *k*-mer matching-based approach, which relies on exact matches between single-copy, unique 15-mer sequences derived from each contig and the male or female reads. Ultimately, we calculated the percentage of these unique, single-copy 15-mers in each contig that does not match female reads (Carvalho and Clark 2013).

Our coverage-based analysis of Y-derived sequences in the three species’ assemblies identified many high-confidence Y contigs (Figure 4A). We identified 169, 88 and 61 contigs that are putatively derived from the Y chromosome in *D. americana, D. novamexicana* and *D. virilis*, respectively (File S2). The respective combined lengths of these contigs in each species is 24.1 Mb, 8.9 Mb, and 14.8 Mb (Figure 4B), suggesting that these species might substantially differ in Y chromosome length. The vast majority of Y-derived contigs are ≤500 kb, with only a handful being ≥1 Mb.

**Figure 4.**
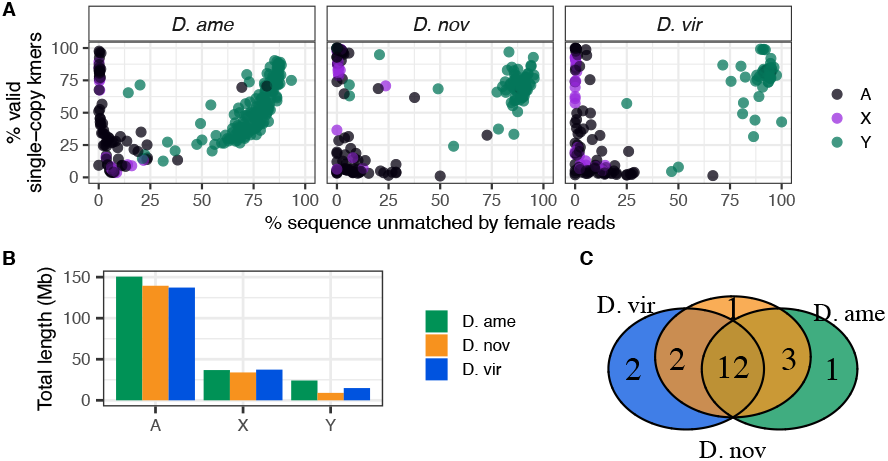
Genomic analysis of Y chromosome-derived contigs. (A) Genomic contigs plotted based on *k*-mer matches of female-derived reads (*x*-axis) and the percentage of single-copy *k*-mers (*y*-axis). Each point is colored based on coverage ratio between male and female mappings. Based on k-mer analysis, Y contigs will be skewed toward the right (largely unmatched by female reads), and high confidence Y contigs should have a high percentage of single-copy k-mers. (B) Aggregate length of genomic contigs on the autosomes, X and Y chromosome for each species. (C) Number of shared and unique Y-linked genes among the three species.

To examine the gene content on these putative Y contigs we used the *D. novamexicana* gene annotations from NCBI and mapped those annotations onto the other two species’ genomes. We ultimately identified 21 single-copy protein coding genes on the Y chromosomes of these species, 12 of which are identified in all three species and between 1-3 that are unique to at least one of the species (Figure 4C, Table S2). Nearly half of the functionally annotated genes on the validated Y contigs in each species codes for components of the dynein complex, which is an essential component of the sperm tail axoneme in *Drosophila*.

### Transposable element composition in the D. virilis group

Our high-quality PacBio assemblies resulted in improved TE annotation compared to the earlier *D. virilis* assembly, especially for retroelements. We identified and annotated TEs with RepeatModeler2 and RepeatMasker on both the Caf1 genome (produced from Sanger sequencing in 2007; Clark *et al*. (2007)) and our PacBio assembly. The Caf1 assembly had a higher proportion of unknown (thus likely incomplete) elements, and a smaller proportion of LTR and non-LTR retrotransposons (Figure S3).

The *D. virilis* group species genomes are rich in repetitive sequences. In addition to the simple satellites that compose approximately 40% of the genome—which are largely excluded from assemblies (Flynn *et al*. 2020b), the genomes also contain abundant transposable elements and other interspersed complex repeats. The total percentage of the genome assembly that is composed of interspersed repeats is similarly high in *D. virilis* and *D. americana* (∼30%), and is lower in *D. novamexicana* (22%) (Figure 5A). Notably, *D. virilis* has higher amounts of retrotransposons—especially LTRs—than both *D. novamexicana* and *D. americana*. The model species *D. melanogaster* has a similar proportion of its genome made up of its TEs (Flynn *et al*. 2020a) as species in the virilis group. However, *D. melanogaster* has less than 1% of its genome composed of Helitron-derived elements and does not contain the interspersed tandem repeats we will describe here. The inferred Y chromosome assembly of each virilis group species was composed of approximately 90% TEs (Figure 5C). The composition of special types of repetitive elements, such as DINE-tandem repeats and interspersed tandem repeats, are described in the sections that follow.

**Figure 5.**
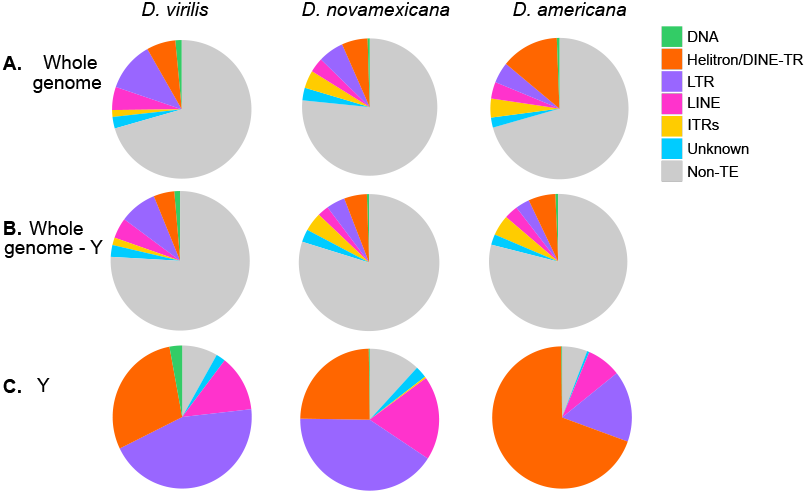
Transposable element and interspersed repeat composition by subclass. The genomes were masked with the modified RepeatModeler2 libraries using Repeat-Masker, and parsed using parseRM. (A) Whole genomes; (B) Whole genomes except for contigs called as the Y chromosome; (C) Contigs called as the Y chromosome only.

The sequence divergence of each genomic copy of a given TE to its respective consensus sequence is often used as a proxy of the age of the insertion. TE copies with low divergence from the consensus indicate that these TE instances represent recent transpositions in the genome. A peak of high abundance TE copies that are each highly diverged (e.g., over 10%) from their respective consensus often represents an ancient expansion of elements that may no longer be transpositionally active. We generated landscape plots (percent divergence vs. abundance) to demonstrate the relative age structure of different TE subclasses (Figure 6A). All species contained both recently- and anciently-active retrotransposons (LINEs and LTR elements), with *D. virilis* having the highest fraction of the genome (17% compared to 9% in *D. americana* and *D. novamexicana*). All species contained only a small fraction (0.5-1%) of DNA elements (with terminal inverted repeats or TIRs). We also compared TE composition across 100 kb windows of the three genomes (Figure 6B, Figure S5). Overall, *D. virilis* has more euchromatic copies of LTR elements. As expected, the density of TEs increases toward the pericentromeric heterochromatin of the chromosome arms and close to where the assembly ends, after which several Mb arrays of simple satellites begin.

**Figure 6.**
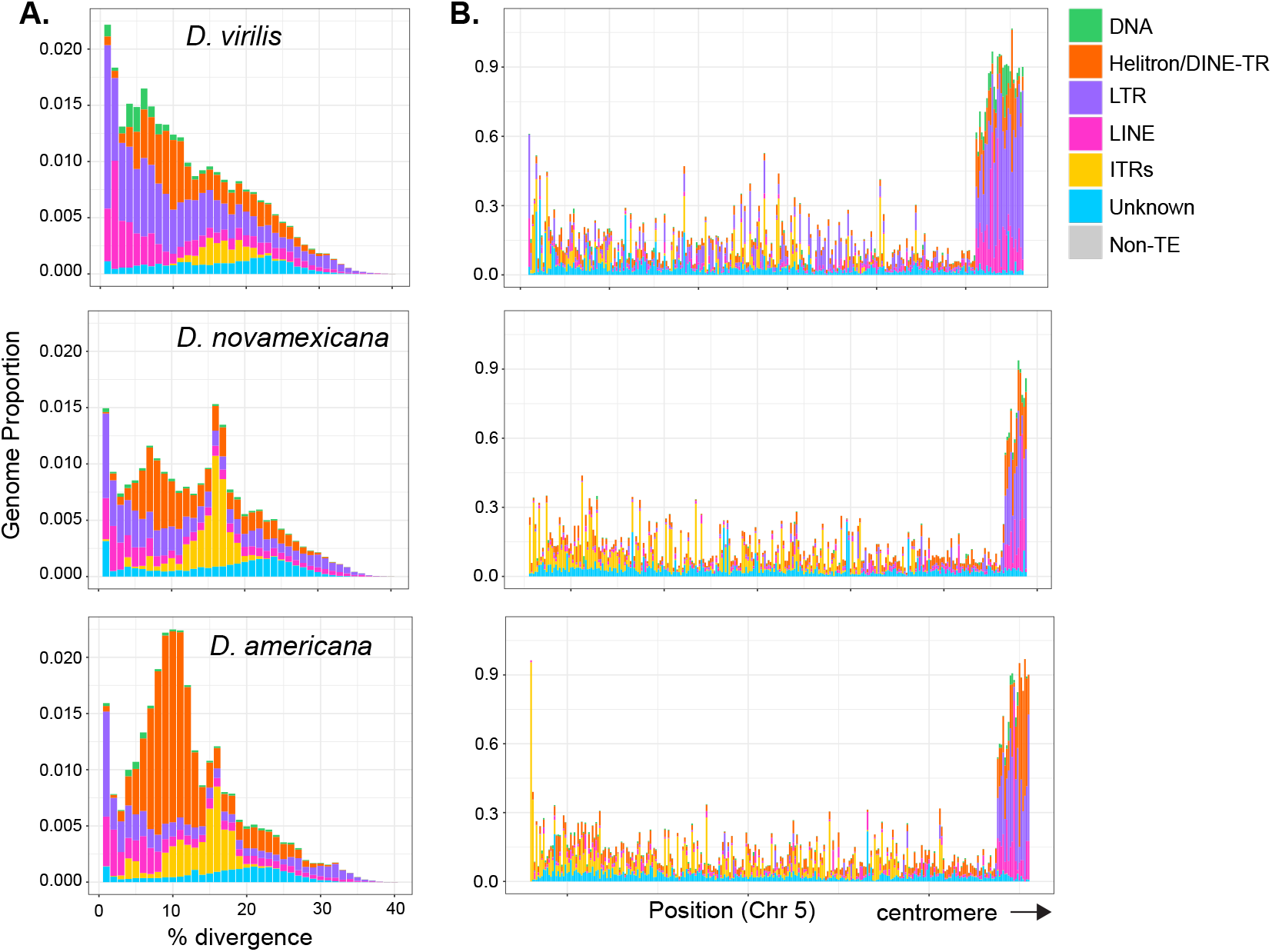
Interspersed repeat landscapes and density in the virilis clade. (A) Genomic proportions of each repeat subclass in 1% divergence bins compared to the representative sequence from each genomic copy’s respective subfamily. TEs with recent insertions have low divergence values, and older insertions have higher divergence values. The divergence of DINE-TRs and ITRs (orange and yellow) may not necessarily reflect their age since their mechanism of propagation is not understood. (B) TE density in 100 kb windows by subclass across a representative chromosome, Chr 5 (left to right; telomere to centromere-proximal).

### DINE-tandem repeats and Interspersed tandem repeats have distinct patterns of evolution

We found DINE-TRs to be highly abundant, and especially so in *D. americana*, with 13.5% of the genome composed of DINE-TRs or Helitron-derived repeats, compared to about half of that in *D. virilis* and *D. novamexicana* (Figure 4A). DINE-TRs had similar abundance among all three species on the autosomes and X chromosome (5-6%, Figure 5B), but the Y chromosome in *D. americana* experienced a massive increase in DINE-TR copy number. DINE-TRs make up 69% of the Y in *D. americana* but only 25% of the Y in *D. novamexicana* (Figure 5C). Therefore, the DINE-TR expansion in *D. americana* is attributed to the Y chromosome-specific DINE-TR expansion. All species’ Y chromosomes contained some “young” insertions of retrotransposons and a variable amount of degrading retrotransposons (Figure S6). *D. americana* contained very few degrading copies of retroelements with >5% divergence from the consensus, suggesting almost complete takeover of inactive elements by the DINE-TR repeats. “Takeover” of the Y chromosome by Helitron-related elements has also been found in *D. pseudoobscura* and in *D. affinis* (Nguyen *et al*. 2022).

Another distinct type of repeats in the virilis group are what we call ITRs (interspersed tandem repeats), which are represented by the previously-discovered pvB370 and 172TR elements and their variants (Heikkinen *et al*. 1995; Abdurashitov *et al*. 2013). We find that these repeats are found in medium length tandem arrays (∼0.5 – 10 kb) that are abundantly dispersed in ∼2000 loci in the genome (Table S3). Our long-read assemblies revealed that 172TR and pvB370 and their variants (ITRs) are especially abundant in the *D. novamexicana* and *D. americana* genomes (∼4%, compared to 1.6% in *D. virilis*, Figure 5A), mostly interspersed in the euchromatic chromosome arms, for example, on Chr 5 (Figure 6B). However, they are absent from the Y chromosome (Figure 5C, Figure S6). The distribution of ITRs on euchromatic arms of *D. novamexicana* and *D. americana* mirror the distribution of LTR elements in *D. virilis* (Figure 6B).

This distribution is in stark contrast to that of typical satellite DNAs present in the pericentromeric heterochromatin of *D. virilis* (Flynn *et al*. 2020b). The mean length of ITR loci is 5.5 kb in *D. americana*. How ITR elements increase their copy number in the genome is unknown, as they do not contain coding sequences or other features of TEs with known mobilization mechanisms. We found that the higher abundance of ITRs in *D. novamexicana* and *D. americana* genomes is associated with longer tandem arrays rather than more loci (Table S3), suggesting that satellite-like evolution dominates the abundance changes we found. However, we found that these ITRs are also capable of mobilizing to new locations in the genome. We searched conservatively for *de novo* insertions of ITR elements present in one *D. americana* strain but absent in the other two. We confidently identified a 1.5 kb novel insertion of 172TR on Chromosome 5 (in an intron of gene *LOC6624736*) in *D. americana* G96 but not in ML97.5 or SB0206, where the continuous and unique insertion site is intact (Figure 7). This suggests that 172TR elements are capable of mobilizing to new loci in the genome, further blurring the distinction between transposable elements and satellite sequences (Meštrović *et al*. 2015; Zattera and Bruschi 2022).

**Figure 7.**
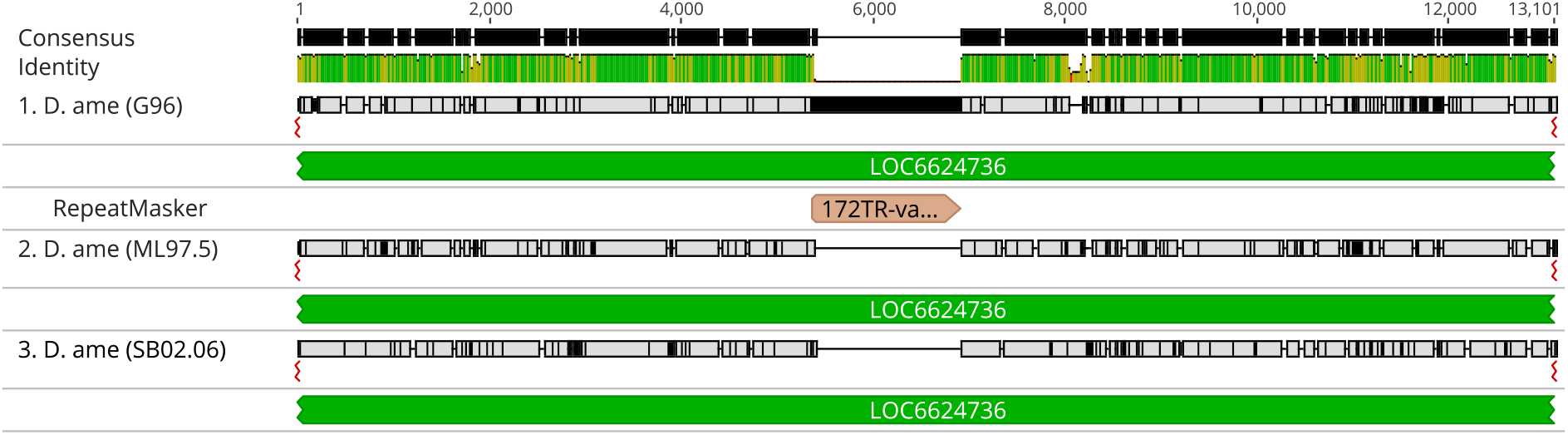
Genome alignment of the three *D. americana* strains analyzed demonstrates a novel insertion of a 1.5 kb array of the ITR element 172TR into chromosome 5 in G96. The insertion is located within an intron of the gene LOC6624736.

Both DINE-TRs and ITRs demonstrate discrete peaks in the landscape histogram plot, with DINE-TRs peaking at 8-10% and ITRs peaking at 15-17% divergence (Figure 6A). For typical transposons, peaks at high divergence would reflect an ancient burst of activity. In this case, it is possible that there was an ancient burst followed by differential levels of deletion in species of the virilis group. Alternatively, sequence diversification could occur during mobilization. The polymorphic (and thus recent) insertion array of 172 TR has a lower sequence divergence than the most common divergence level (3% compared to 15-17%). Further research is required to determine the mechanisms of copy number change, sequence diversification, and mobilization of DINE-TRs and ITRs.

### Novel elements unique to single species’ genomes

Little is known about the birth rate of new TEs in a genome, which may form *de novo* or may be introduced through horizontal transfer. Here, we searched conservatively for TEs that were novel to each genome and not present in the other two species’ genomes. We only included elements with recent copies to exclude false positives caused by old differentially diverged copies (details in methods). We found 2, 1, and 1 novel elements in our assemblies of *D. virilis, D. novamexicana*, and *D. americana*, respectively (Table 2). The *D. virilis* novel element rnd-1_family-95, which was incorrectly classified as a Gypsy element by RepeatModeler2, is present as a complex tandem array on two unplaced (likely heterochromatic) contigs, which were not identified as being on the X or Y chromosomes. To our knowledge, this putatively species-specific tandem repeat has not been identified in previous studies that used both short- and long-read sequencing to characterize repeats (Flynn *et al*. 2020a; Silva *et al*. 2019). It totals 313 kb and is likely present in one or two arrays. Short-read data from other *D. virilis* strains (described in this manuscript) map to rnd-1_family-95, indicating the repeat is not unique to the genome assembly strain.

**Table 2.** Novel repeats or transposable elements found to be unique to each species.

| Species | Novel element name | Novel element location(s) | Consensus length | Notes |
| --- | --- | --- | --- | --- |
| *D. virilis* | rnd-1_family-95# LTR/Gypsy | Unplaced contigs *tig00001890*,<br>*tig00002154* | 11505 bp<br>(mix of 1132 bp and<br>820 bp monomers) | Structured as a complex tandem repeat<br>(the 820 bp monomer has an internal deletion<br>compared to the 1132 bp monomer); homology<br>to Gypsy is minimal. |
| *D. virilis* | rnd-1_family-144#LINE/I-Jockey | Unplaced contigs *tig00001890*,<br>*tig00001978*, *tig00002154*,<br>*tig00002244*, *tig00003829*,<br>*tig00003830* | 4834 bp | Highly similar to Repbase sequence HeT-A3_DVi,<br>which covers 84% of the sequence at 99% identity.<br>A gag domain is identifiable. |
| *D. novamexicana* | rnd-1_family-329#Unknown | Present on all Muller elements<br>and a few unplaced contigs | 346 bp | No conserved domains identifiable, no TSD<br>identified from REPCLASS. |
| *D. americana* | rnd-1_family-159#Unknown | Y contig *tig00002773* | 767 bp | Likely the same element as below: overlaps 77 bp<br>with below sequence and found in tandem with it. |
| *D. americana* | rnd-1_family-109#DNA/hAT-Ac | Y contig *tig00002773* | 421 bp |  |

The second novel element in *D. virilis*, rnd-1_family-144, is a Jockey element in the same family as the Repbase-annotated HeT-A3_DVi sequence (99% similarity across 84% of the sequence) and contains an identifiable gag domain. This element is found on multiple unplaced contigs which are likely heterochromatic but not in tandem with other Jockey elements as may be expected of telomere-specific HeT-A elements. In *D. novamexicana*, the novel element rnd-1_family-329, is present in an interspersed manner across all major chromosome arms in addition to some unplaced contigs. We were not able to identify any classifiable aspects of this element as it had no homology to known proteins or other TEs, and no target site duplications (TSDs) were identified using the TSD module of the REPCLASS software (Feschotte *et al*. 2009). Therefore, by elimination, a possible classification is a non-autonomous Helitron.

Finally, in *D. americana*, we identified two elements (rnd-1_family-159 and rnd-1_family-109) but they are likely a single element based on their overlap in sequence identity and tandem distribution. This element has homology to a DNA (TIR) element and seems to be Y-specific, being present only on a single Y contig. The *D. americana* novel Y-specific element is also present in the same distribution pattern in the SB02.06 and ML97.5 genomes.

## Conclusions

The genomes presented in this work represent a major advance in the comparative genomics resource for a set of closely related, experimentally tractable *Drosophila* species. Coupled with the available genome annotations and those produced in this study, the virilis group is now poised to become a *bona fide* genetic model system. Here we highlight the main attributes of these genome assemblies and confirm major structural rearrangements differences between species. We also confidently identify Y-derived contigs and Y-linked protein-coding genes. Our work has also uncovered evolutionary patterns of poorly understood repetitive elements. We found that *D. americana* experienced a proliferation of DINE-TRs specifically on the Y chromosome, resulting in two-thirds of the chromosome being composed of this repeat. This Y chromosome-specific expansion represents the greatest magnitude of sequence-level divergence between sister species *D. americana* and *D. novamexicana*. Furthermore, we found a high abundance of peculiar interspersed tandem repeats (ITRs) across the virilis species group. ITRs have unknown mechanisms of expansion and mobilization, but we found one instance of a polymorphic insertion within *D. americana*, demonstrating that these elements are capable of transposition. Furthermore, we found putatively species-specific repeats in each genome, all with different distribution patterns (heterochromatic in tandem, Y-specific, interspersed). Further studies may identify the origins of species-specific repeats and their evolutionary significance.

## Materials an Methods

### PacBio long-read sequencing

High molecular weight (HMW) DNA was extracted from inbred males, using an in-house method. Briefly, flash frozen flies were prepared by putting 50 males into a 1.5 ml microfuge tube and quickly frozen in liquid nitrogen and stored at -80°C. For extraction, tubes of flies were put on ice and 500 ul of room temperature buffer FLY-A was immediately added; (Fly-A: Tris HCl buffer 0.1 M pH 8.0, EDTA 0.1 M pH8, SDS 1%). Flies were homogenized with a nylon fitted sterile pestle with 5 twist motions. Tubes were Incubated at 60°C for 30 min, then at 37°C for 5 min. 2.5 µl of proteinase K solution (20 mg/ml in TE) was added to each tube followed by gentle mixing. Tubes were incubated for 30 min at 37°C followed by addition of 70 µl of 4M potassium acetate and gentle mixing and incubation on ice for 20 min. After centrifugation at 4°C for 30 min at 13,000 rpm, a wide bore tip was used to remove supernatant to a fresh tube. An equal volume of chloroform/isoamyl alcohol (24:1) was added followed by gentle rocking of tubes by hand forty times. Tubes were centrifuged for 5 min at 4°C and 10,000 rpm. The upper phase was then pipetted to a fresh tube that contained 350 µl of isopropanol. The phases were mixed by gentle rocking at which time threads of DNA were observed. DNA was collected by centrifugation, washed with 70% ethanol, dried and dissolved in 100 µl of TE. RNA was removed with the addition of 3 ul RNaseA solution and incubation at 37°C for 30 min. DNA quality was checked by Nanodrop for purity, CHEF gel (Bio-Rad) for size, and Qubit for total mass. Sequencing libraries were constructed according to Pacbio standard methods and final size selections on Blue Pippin (Sage Sci) using the S1 marker. Pacbio sequencing was performed on a Sequel instrument (chemistry version 2.0) following the manufacturer’s standard methods. Sequencing was collected for 10 hr per SMRT cell to approximately 100X genome coverage. Strains used were as follows: *D. virilis*: 15010-1051.87, *D. novamexicana*: 15010-1031.14, *D. americana*: G96. Raw reads are available under NCBI SRA accession PRJNA475270.

### Nanopore long-read sequencing

HMW DNA from two *D. americana* strains (SB02.06 and ML97.5) was prepared using the approach recommended by Oxford Nanopore Tech with slight modifications. Briefly, ∼200 flash-frozen male flies were homogenized in a nuclear isolation buffer (0.35M sucrose, 0.1M EDTA, 50mM Tris-HCl) using a TissueRuptor II with 2 x 15 second pulses on speed 2. Fly homogenate was filtered through 1 layer of 200 µm nylon mesh and centrifuged at 3500g for 15 minutes at 4°C. The supernatant was then discarded and the pellet was gently resuspended in 5 ml Buffer G2 and 95 µl proteinase K (QIAGEN Blood and Cell Culture DNA Midi Kit). The lysate was then incubated at 50°C for 45 min with gentle mixing at 100 rpm. The lysate was then poured into an equilibrated QIAGEN Genomic-tip 100/G column and purified according to manufacturer’s instructions (QIAGEN). HMW DNA was eluted overnight at room temperature in 150 µl TE buffer (10 mM Tris-HCl, 1 mM EDTA, pH 8.0) with gentle shaking at 300 rpm. The DNA was quantified on a Qubit fluorometer and fragment size distribution was checked on an Agilent TapeStation 4200 using the Genomic DNA ScreenTape assay. Sequencing libraries were prepared using the Genomic DNA by Ligation kit (SQK-LSK 109) according to manufacturer instructions with 1 µg of input DNA. Two R9 flow cells were used for each *D. americana* strain, with a final read output of 1.3-1.6 million reads.

### Illumina short-read sequencing

Several strains were sequenced with Illumina low or medium coverage sequencing (see Table S1 for list of strains). For these libraries, we collected ∼5 adult male flies for DNA extraction with Qiagen DNeasy blood and tissue kit and prepared with Illumina PCR-free library prep. Libraries were run on Illumina NextSeq 500 with 1x150 bp reads. Strains Toyama15, vzzp01, vww8, England1430, England1431 were sequenced from fly DNA extracted previously (kindly provided by Anneli Hoikkala, University of Jyväskylä). Toyama15, vww8, and England 1431 were male while vzzp01 and England1430 were female.

### Assembly

For the *D. virilis* and *D. novamexicana* PacBio genomes, contig assembly was performed with MECAT v1.3 (Xiao *et al*. 2017) and polished with Pilon v.1.18 (Walker *et al*. 2014). For the *D. americana* G96 PacBio genome, Canu Koren *et al*. (2017) (genome size parameter set to 275Mb) and Falcon (Chin *et al*. 2016) assemblies were used for scaffolding with Quickmerge (see below). Nanopore long-reads were assembled using Canu (v.2.1.1) with the genome size parameter set to 200Mb.

### Scaffolding

To generate Muller element scaffolds from the raw assembly, the *D. novamexicana* and *D. virilis* MECAT polished assemblies were aligned to the dvir1.06 scaffolded genome assembly with NUCmer (Kurtz *et al*. 2004; Schaeffer *et al*. 2008; Ahmed-Braimah and Sweigart 2015). The *D. americana* raw assembly was first scaffolded using QuickMerge (Chakraborty *et al*. 2016) using the Canu and Falcon assemblies as input, and subsequently scaffolded by aligning to the dvir1.06 scaffolded genome with NUCmer. The *D. americana* assemblies contained duplicate contigs for a number of locations along the Muller elements, and thus duplicate contigs were manually inspected and removed from the assembly of placed Muller Elements (unplaced contigs were left in the assembly). For all three species’ assemblies, the arrangement of contigs along the Muller elements was inspected manually using dotplot alignments and the order of contigs was recorded (File S3). (The chromosomal inversion differences between *D. americana*/*D. novamexicana* and *D. virilis* were taken into account when analyzing the dot-plots). Contig sequences for each Muller element were then concatenated (without gaps) to generate the Mullerized, scaffolded assemblies.

### Transposable element discovery

Transposable elements were annotated *de novo* in the three PacBio assemblies using RepeatModeler2 (Flynn *et al*. 2020a). RepeatModeler2 uses a genome assembly to produce consensus sequences for genome-wide transposable element families. RepeatModeler2 has a specific module to build full-length consensus sequences for LTR elements, which are abundant in *Drosophila*. We used the Muller element-scaffolded PacBio genomes as input into RepeatModeler2 and used options -LTRStruct -LTRMaxSeqLen 10000. *D. virilis* group species contain a diversity of complex tandem repeats and interspersed-and-tandem repeats (ITRs) that are detected by RepeatModeler2. DINE-tandem repeats (DINE-TRs) have evolved from an internal expansion within the DINE element (Dias *et al*. 2015), and we found related repeats to be diverse and encompassed by several RepeatModeler2 families. ITRs like pvB370 and 172TR also had multiple families but were classified as “Unknown”. We used BLAST to identify RepeatModeler2 sequences that have significant similarity to the consensus sequences of pvB370 and 172TR repeats to curate and clean the library. We then removed the sequence with similarity to the consensus sequence or re-classified it appropriately. DINE-TRs were the most abundant and diverse type of complex tandem repeats. It was not possible to easily differentiate between DINE-TRs and other Helitron-derived repeats, thus we grouped Helitrons and DINE-TRs together. We used RepeatMasker to annotate the genome with the library produced from RepeatModeler2 that we edited as described above. We then used ParseRM (https://github.com/4ureliek/Parsing-RepeatMasker-Outputs/blob/master/parseRM.pl) to process the RepeatMasker output for genome proportions and percent divergence landscape proportions. All pie charts and landscape plots were produced in R using ggplot2. We also used the RepeatMasker output along with custom bash scripts to visualize the density of TE subclasses in 100 kb windows across each chromosome arm in each species. All analysis scripts and the TE libraries for each species can be accessed on a GitHub repository.

We found TE families that were unique to each genome by first clustering the TE libraries with CDHIT (Li and Godzik 2006). Next we BLASTed the sequences that did not cluster with any other sequences (singleton elements) against the other two genomes. Only singleton elements that contained no significant hits (≥100 bp at ≤20% sequence divergence) to both other genomes were considered truly unique. Further we narrowed down to singleton elements with at least 5 kb in the genome and with average sequence divergence <10%, to remove old inactive TE copies that have differentially diverged across species.

In the three different *D. americana* genomes, we searched for confident polymorphic insertions of ITR elements; i.e. insertions that were in one of the genomes but not the other two. First, we focused only on insertions in one genome at a time that were not flanked in a 1 kb region by other annotated repeats. Next, we extracted the left and right flanking 100 bp sequences of each eligible ITR locus. We merged the flanking sequences to a 200 bp continuous sequence, which would represent an intact insertion site. We then blasted the intact insertion site against the other two genomes. We required a unique and continuous blast hit with at least 90% identity over the 200 bp. We manually checked putative polymorphic insertions with multiple sequence alignments.

### Structural rearrangement analysis

The scaffolded genome assemblies for the three species were aligned using NUCmer to identify the precise locations of rearrangement breakpoints between species. First, we omitted initial NUCmer alignment matches that were smaller than 100 bp. We then filtered out alignments that were below 90% sequence identity and spanned less than 1000 bp in either the query or reference sequence. Finally, we examined the coordinates of each segment match between query and reference and identified contiguous tandem matches along the five Muller elements that were either inverted or translocated. The boundaries of these inverted matches were deemed as rearrangement breakpoints (File S1).

To find repetitive elements in the breakpoint regions, we used bedtools to intersect the predicted breakpoints with the TE annotation in each species. If TEs were present, the regions were extended to the TE’s called end-point, normalizing each TE-intersected breakpoint to 100%. Within the extended regions, we calculated the percentage of each TE subclass and DAIBAM DNA elements, which we expected to be enriched based on previous work (Reis *et al*. 2018). This enrichment was especially evident in *D. novamexicana*, thus we performed a permutation test where we randomly selected regions of the genome of the same size as the predicted breakpoints, and calculated the total abundance of DAIBAM in the extended regions, and repeated this 1000 times.

### Divergence and Polymorphism

To assess the genomic pattern of nucleotide diversity we used publicly available short-read genome sequence data from multiple strains of *D. americana, D. novamexicana*, and *D. virilis*. Illumina short reads from each strain were assessed for quality before mapping to the genomes of their respective species using BWA MEM (Li and Durbin 2009) with default parameters. Single nucleotide variants for each strain were called using GATK’s HaplotypeCaller (McKenna *et al*. 2010), and SNP calls were merged for downstream analysis. We measured population-level nucleotide diversity (*p*) and pairwise species divergence (*D*_*xy*_Dxy) across non-overlapping 20 kb sliding windows using pixy, which accounts for invariant sites when calculating these parameters (Korunes and Samuk 2021).

### Identification of Y chromosome contigs

To identify contigs from each of the three assemblies that are derived from the Y chromosome, we performed two levels of analysis: (1) Matching of unique *k*-mers that are derived from either a male or female Illumina paired-end sequencing libraries to our assemblies (Carvalho and Clark 2013), and (2) coverage-based differences between male and female samples. The Illumina samples were based on sequencing that was separately performed in males and females as part of a previous study (see Table S1 for samples description; Ahmed-Braimah et al. 2017), and coverage in each sample was ∼22X for each species/sex.

To perform the *k*-mer matching analysis on the male/female reads, we used the “YGS” software (Carvalho and Clark 2013). Briefly, male and female reads were filtered using the Jellyfish “count” function, and a quality score cutoff of 5 and *k*-mer size of 15. We then created a bit array trace of the filtered *k*-mers using the YGS.pl script for both filtered female and male reads, in addition to bit array traces of the genome contigs. Finally, we scanned each contig for the percentage of unique, single-copy *k*-mers that fail to match female reads. Analysis details can be found in the project’s GitHub repository under the “YGS” folder.

To perform the coverage-based analysis we separately mapped male or female reads from a given species to that respective species’ genome assembly using Bowtie2 with default parameters. We then used the coordinate-sorted alignment BAM file as input for analysis using QualiMap (García-Alcalde *et al*. 2012) to output average coverage statistics for each contig. We validated a subset of high confidence Y contigs in *D. novamexicana* by designing at least one pair of PCR primers that amplify a 200-300 bp non-repetitive region of each contig that had a skewed male:female coverage ratio. PCR reactions were performed on a male and female DNA sample from *D. novamexicana* and examined on a gel. A contig was considered validated if it amplified in the male sample but not female. To perform gene content comparisons between validated Y contigs we mapped the gene annotations from the *D. novamexicana* assembly (GCF_003285875.2) onto *D. americana* and *D. virilis* using Liftoff (Shumate and Salzberg 2020). (We used the *D. novamexicana* annotations because we only validated Y contigs by PCR in this species). To identify the high confidence protein-coding genes in each species’ Y chromosome we first identified the Y contigs that contain validated Y genes in the *D. novamexicana* annotation, then captured all the protein-coding genes on these contigs independently using the lifted annotation.

## Data availability

Short-read sequencing data produced in this paper are available at NCBI SRA PRJNA981214. We also analyzed previously-published data from NCBI SRA PRJNA548201 (details in Table S1). The RS2 non-scaffolded assembly for *D. virilis* is available at https://www.ncbi.nlm.nih.gov/assembly/GCF_003285735.1. The RS2 non-scaffolded assembly for *D. novamexicana* is available at https://www.ncbi.nlm.nih.gov/assembly/GCF_003285875.2. The PacBio genome assemblies we used in analyses are Muller-element scaffolded assemblies available at NCBI Assembly PRJNA982537. The two *D. americana* assemblies, ML97.5 and SB02.06, have project accessions PRJNA991588 and PRJNA923584, respectively. The raw PacBio reads are available at PRJNA475270. All scripts to reproduce our analyses are located in the project’s GitHub repository.

## Acknowledgements

We thank Jianwei Zhang for assembling the genomes produced by PacBio sequencing. We also thank Cedric Feschotte, Daniel Barbash, and members of the Clark lab for providing advice on the manuscript. This work was supported by NIH grant R01GM116113 to RAW, ML and AGC and R35GM147454 to YHAB.

