## Supplementary material for "High quality genome assemblies reveal evolutionary dynamics of repetitive DNA and structural rearrangements in the *Drosophila virilis* sub-group": Table S1, Table S2, Table S3, Figure S1, Figure S2, Figure S3, Figure S4, Figure S5, Figure S6

**Table S1** Strains sequenced in this study.

| Strain number | Shortform | Species | Accession |
|---|---|---|---|
| *15010-1051.51* | vir51 | *D. virilis* | PRJNA548201 |
| *15010-1051.52* | vir52 | *D. virilis* | PRJNA548201 |
| *15010-1051.86* | vir86 | *D. virilis* | PRJNA548201 |
| *15010-1051.47* | vir47 | *D. virilis* | PRJNA548201 |
| *15010-1051.49* | vir49 | *D. virilis* | PRJNA548201 |
| *15010-1051.85* | vir85 | *D. virilis* | PRJNA548201 |
| *15010-1051.08* | vir08 | *D. virilis* | PRJNA548201 |
| *15010-1051.00* | vir00 | *D. virilis* | PRJNA548201 |
| *15010-1051.118* | vir118 | *D. virilis* | PRJNA548201 |
| 15010-1051.48 | vir48 | *D. virilis* | PRJNA548201 |
| 15010-1051.87 | vir87/GDvir* | *D. virilis* | PRJNA548201 |
| Toyama15 | | *D. virilis* | This study, PRJNA981214 |
| Vzzp01 (f) | | *D. virilis* | This study, PRJNA981214 |
| vww8 | | *D. virilis* | This study, PRJNA981214 |
| England1430 (f) | | *D. virilis* | This study, PRJNA981214 |
| England1431 | | *D. virilis* | This study, PRJNA981214 |
| amMK1012 | | *D. americana* | PRJNA548201 |
| amCI0518 | | *D. americana* | PRJNA548201 |
| amCI0515 | | *D. americana* | PRJNA548201 |
| amG96* | | *D. americana* | PRJNA548201 |
| amCI0538 | | *D. americana* | PRJNA548201 |
| amMK0738 | | *D. americana* | PRJNA548201 |
| amSB | | *D. americana* | PRJNA548201 |
| amML975 | | *D. americana* | PRJNA548201 |
| 15010-1031.14 | nov14/GDnov* | *D. novamexicana* | PRJNA548201 |
| 15010-1031.04 | nov4 | *D. novamexicana* | PRJNA548201 |
| 15010-1031.08 | nov8 | *D. novamexicana* | PRJNA548201 |
| 15010-1031.12 | nov12 | *D. novamexicana* | PRJNA548201 |
| 15010-1031.13 | nov13 | *D. novamexicana* | PRJNA548201 |

* Genome strains sequenced with PacBio and assembled.

**Table S2** High confidence Y protein coding genes that with functional annotation.

| species | seqid | gene | product |
|---|---|---|---|
| *D. ame* | tig00000523 | *MSTRG.16118* | Dynein heavy chain 2, axonemal |
| *D. ame* | tig00000545 | *MSTRG.16125* | Transcription-associated protein 1 |
| *D. ame* | tig00000640 | *MSTRG.16192* | Dynein heavy chain 5, axonemal |
| *D. ame* | tig00000640 | *MSTRG.16193* | Retrovirus-related Pol polyprotein from transposon 412 |
| *D. ame* | tig00000656 | *MSTRG.16208* | Retrovirus-related Gag polyprotein from transposon gypsy |
| *D. ame* | tig00000656 | *MSTRG.16217* | Dynein heavy chain 2, axonemal |
| *D. ame* | tig00000686 | *MSTRG.16222* | Dynein regulatory complex subunit 3 |
| *D. ame* | tig00003077 | *MSTRG.16623* | Dynein heavy chain 8, axonemal |
| *D. nov* | NW_020824857.1 | *LOC115767635* | dynein beta chain, ciliary-like |
| *D. nov* | NW_020824875.1 | *LOC115768126* | dynein beta chain, ciliary |
| *D. nov* | NW_020824893.1 | *LOC115768466* | protein Hook homolog 3-like |
| *D. nov* | NW_020824893.1 | *LOC115768469* | protein Hook homolog 3-like |
| *D. nov* | NW_020824896.1 | *LOC115768531* | dynein heavy chain 5, axonemal-like |
| *D. nov* | NW_020824896.1 | *LOC115768532* | dynein heavy chain 5, axonemal-like |
| *D. nov* | NW_020824912.1 | *LOC115768695* | dynein heavy chain 2, axonemal-like |
| *D. nov* | NW_020824944.1 | *LOC115768949* | DBF4-type zinc finger-containing protein 2 homolog |
| *D. nov* | NW_020824946.1 | *LOC115768961* | histone H2A-like |
| *D. nov* | NW_020824946.1 | *LOC115768962* | dynein heavy chain 2, axonemal-like |
| *D. nov* | NW_020824970.1 | *LOC115769090* | dynein heavy chain 8, axonemal-like |
| *D. nov* | NW_020824971.1 | *LOC115769091* | cytosolic non-specific dipeptidase-like |
| *D. nov* | NW_020824975.1 | *LOC115769109* | cilia- and flagella-associated protein 58-like |
| *D. nov* | NW_020824996.1 | *LOC115769151* | dynein regulatory complex subunit 3-like |
| *D. vir* | NW_022587449.1 | *LOC116650070* | transmembrane protein 19-like |
| *D. vir* | NW_022587439.1 | *LOC26530771* | dynein beta chain, ciliary |
| *D. vir* | NW_022587438.1 | *LOC26530795* | dynein heavy chain 8, axonemal |
| *D. vir* | NW_022587438.1 | *LOC26531380* | dynein heavy chain 5, axonemal |
| *D. vir* | NW_022587449.1 | *LOC26531539* | dynein heavy chain 8, axonemal |
| *D. vir* | NW_022587459.1 | *LOC26531779* | dynein beta chain, ciliary |
| *D. vir* | NW_022587459.1 | *LOC26531779* | dynein beta chain, ciliary |
| *D. vir* | NW_022587442.1 | *LOC26531848* | histone H1.2-like |
| *D. vir* | NW_022587474.1 | *LOC26531899* | dynein heavy chain 2, axonemal |
| *D. vir* | NW_022587425.1 | *LOC6635996* | cytosolic non-specific dipeptidase |
| *D. vir* | NW_022587442.1 | *LOC6636766* | DBF4-type zinc finger-containing protein 2 homolog |
| *D. vir* | NW_022587441.1 | *LOC6636891* | protein Hook homolog 3 |
| *D. vir* | NW_022587521.1 | *LOC6637063* | dynein heavy chain 2, axonemal |

**Table S3** Summary of number and length of ITR arrays in virilis group genomes

| Species | Chromosome | Number_arrays | Avg_array_length | Average number of arrays per chromosome | Average length of arrays per chromosome | [illegible] |
|---|---|---|---|---|---|---|
| *D. ame* | 2 | 208 | 7357.09 | 342.6 | 5489.27 | |
| *D. ame* | 3 | 342 | 5897.51 | | | |
| *D. ame* | 4 | 537 | 4216.83 | | | |
| *D. ame* | 5 | 310 | 3761.77 | | | |
| *D. ame* | 6 | 26 | 3265.15 | | | |
| *D. ame* | X | 316 | 6213.15 | | | |
| *D. vir* | 2 | 380 | 1666.87 | 370.4 | 1845.538 | |
| *D. vir* | 3 | 297 | 2611.85 | | | |
| *D. vir* | 4 | 366 | 2081.19 | | | |
| *D. vir* | 5 | 303 | 1556.63 | | | |
| *D. vir* | 6 | 0 | 0 | | | |
| *D. vir* | X | 506 | 1311.15 | 316 | 4633.41 | |
| *D. nov* | 2 | 146 | 6541.23 | | | |
| *D. nov* | 3 | 329 | 4969.43 | | | |
| *D. nov* | 4 | 501 | 4039.77 | | | |
| *D. nov* | 5 | 331 | 3572.31 | | | |
| *D. nov* | 6 | 277 | 1169.4 | | | |
| *D. nov* | X | 312 | 4044.31 | | | |

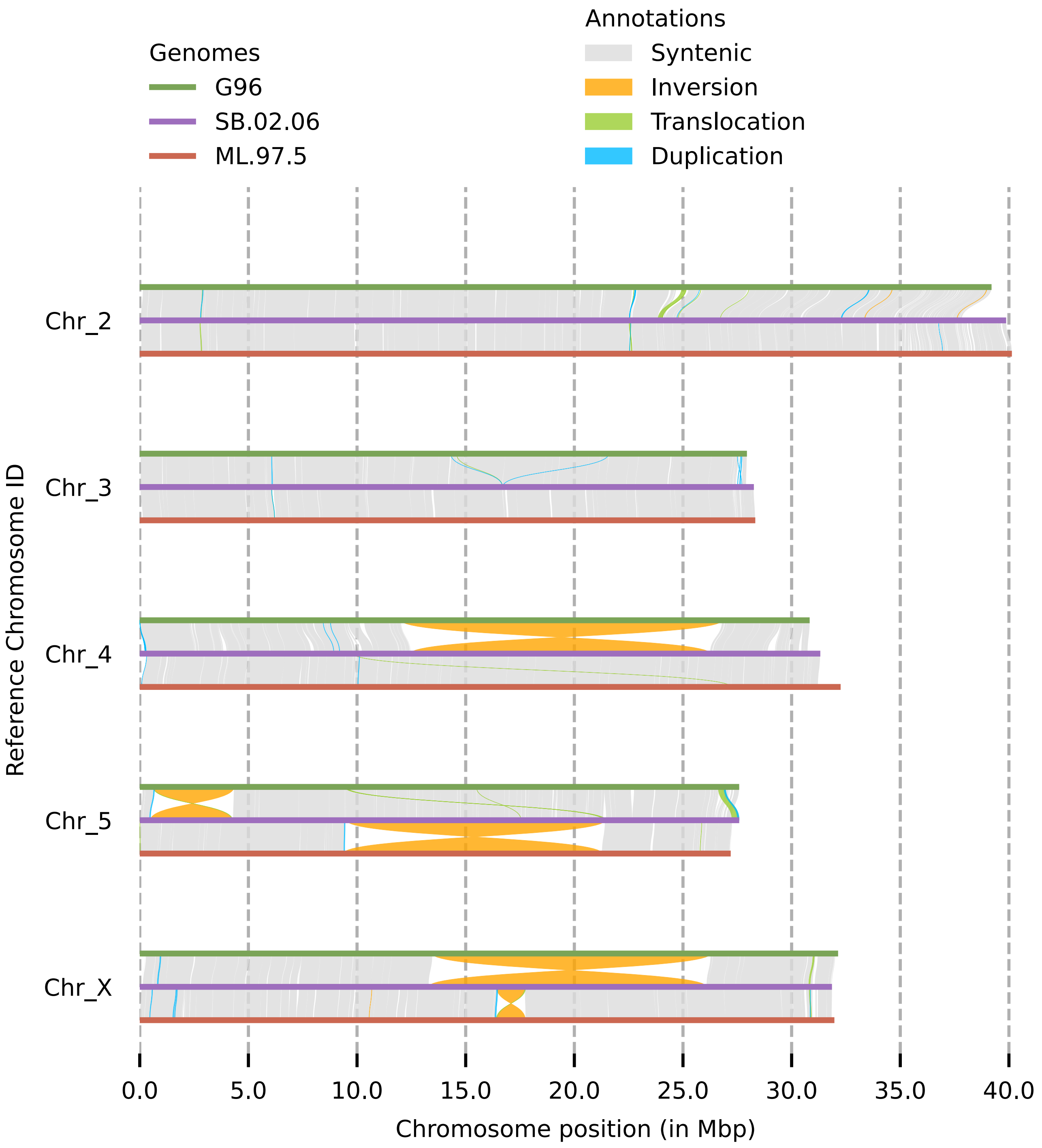


**Figure S1** Whole chromosome alignment of the three *D. americana* genomes showing the main inversion differences between the three strains.

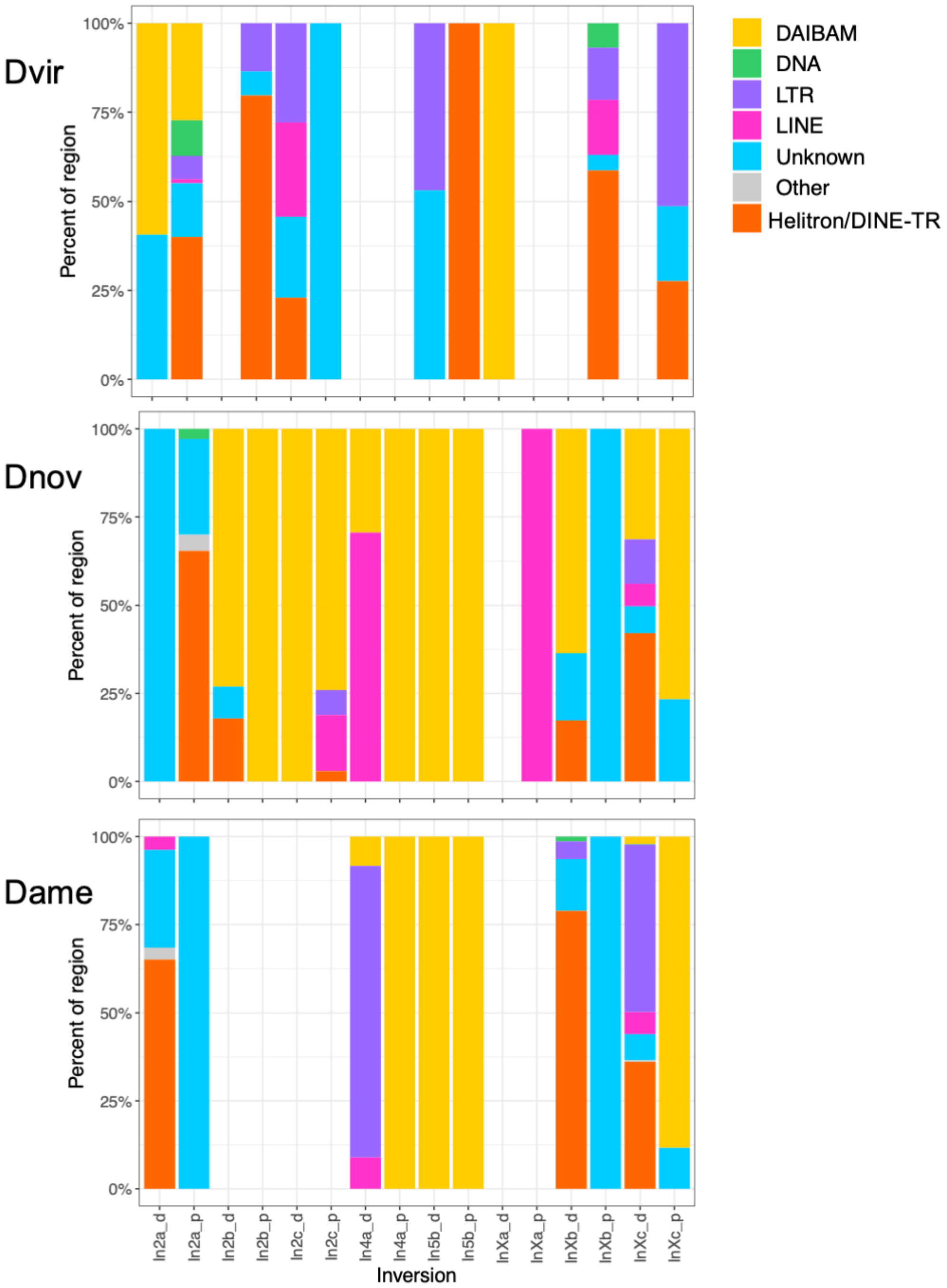


**Figure S2** TE content in regions flanking predicted inversion breakpoints in each species. We intersected the repeat annotation from RepeatMasker with predicted breakpoints. Bars are colored by the TE subclass; bars are absent if there were no repetitive elements in the breakpoint region. If TEs were present, then the regions were extended to the TE's called end point, normalizing each to 100%.

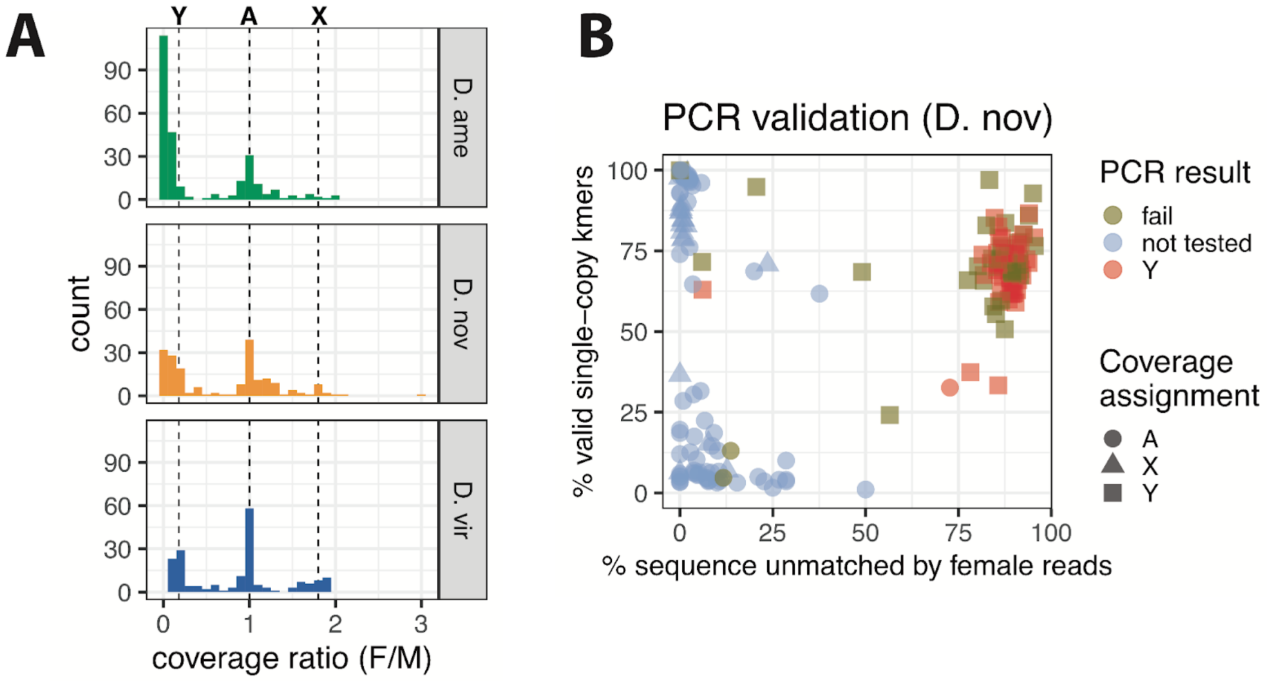


**Figure S3** (A) Contig coverage distribution in the three scaffolded assemblies. The dotted lines indicate the expected coverage ratio for the Y chromosome, autosomes and the X chromosome. (B) Genomic contigs plotted based on *k*-mer matches of female-derived reads (*x*-axis) and the percentage of single-copy *k*-mers (*y*-axis) in *D. novamexicana* with the PCR validation results indicated. Contigs that amplified in the male sample but not the female sample are classified as "Y", whereas those that amplify in both sexes are classified as "failed" to validate.

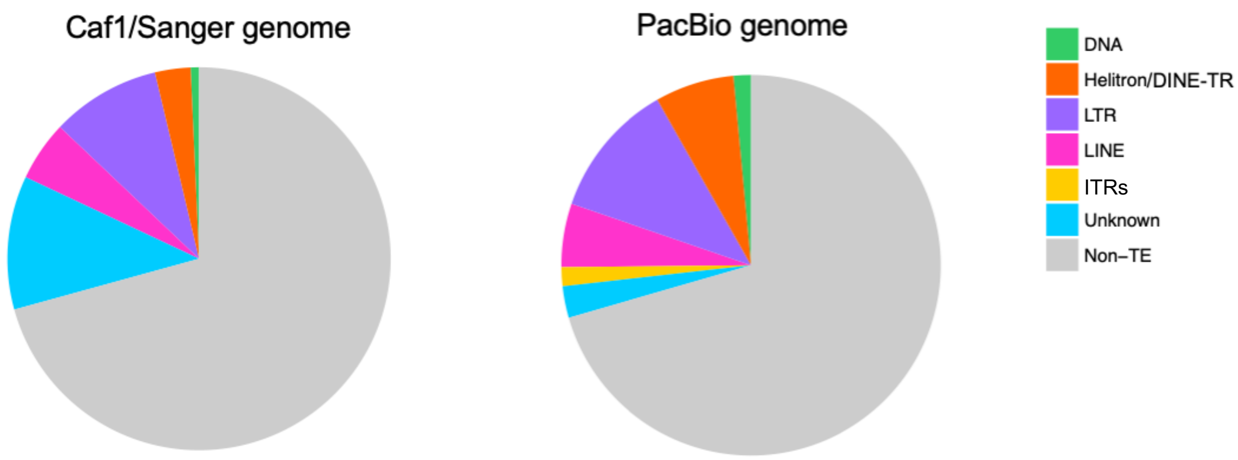


**Figure S4** Transposable element composition in the Caf1/Sanger assembly compared to our new PacBio assembly.

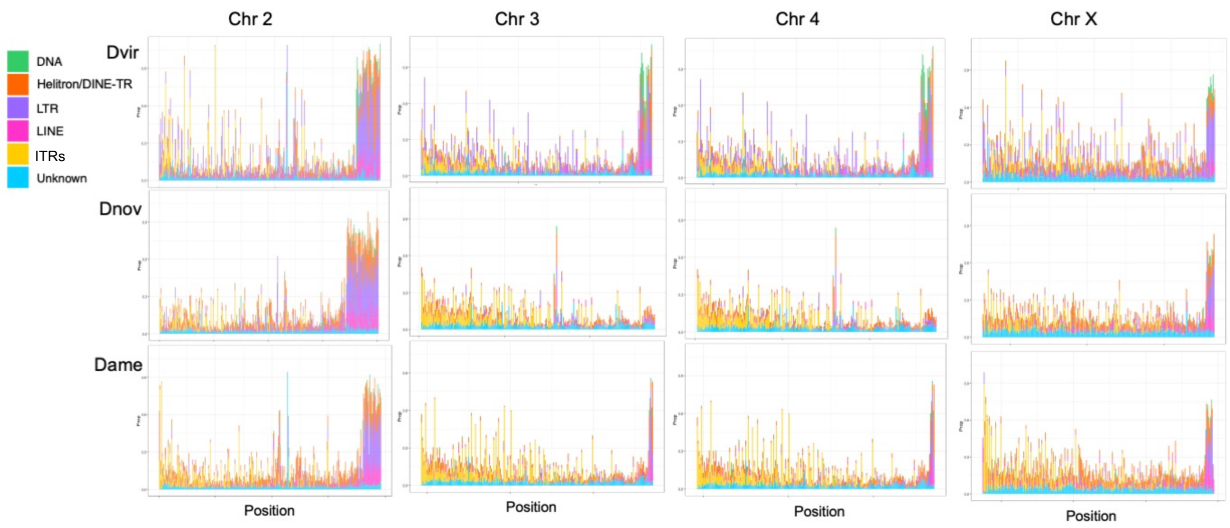


**Figure S5** TE density in by subclass 100 kb windows across chromosomes 2, 3, 4, and X (Chr 5 is included in the main text) in the virilis clade.

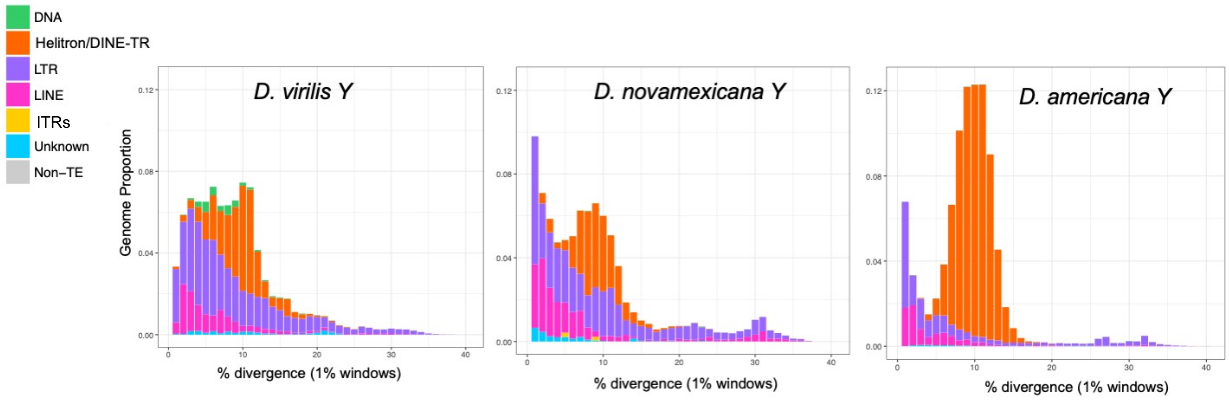


**Figure S6** Divergence landscape of the Y chromosome in *D. virilis*, *D. novamexicana*, and *D. americana*.
